# Differential investment in visual and olfactory brain regions mirrors the sensory needs of a paper wasp social parasite and its host

**DOI:** 10.1101/2021.01.17.427022

**Authors:** Allison N. Rozanski, Alessandro Cini, Taylor E. Lopreto, Kristine M. Gandia, Mark E. Hauber, Rita Cervo, Floria M. K. Uy

**Affiliations:** Department of Biology, University of Miami, Coral Gables, FL 33146, USA; Department of Biology, University of Florence, Sesto Fiorentino (FZ), 50019, Italy; Centre for Biodiversity and Environment Research, University College London, Gower Street, London, WC1E 6BT, UK; Department of Evolution, Ecology and Behavior, University of Illinois at Urbana-Champaign, Urbana-Champaign, IL 61801, USA; Department of Biology, University of Rochester, Rochester, NY 14627, USA

**Keywords:** brain plasticity, coevolution, host exploitation, sensory systems, social parasite, *Polistes dominula*, *Polistes sulcifer*

## Abstract

Obligate social parasites evolve traits to effectively locate and then exploit their hosts, whereas hosts have complex social behavioral repertoires, which include sensory recognition to reject potential conspecific intruders and heterospecific parasites. While social parasite and host behaviors have been studied extensively, less is known about how their sensory systems function to meet their specific selective pressures. Here, we compare investment in visual and olfactory brain regions in the paper wasp *Polistes dominula*, and its obligate social parasite *P. sulcifer*, to explore the link between sensory systems and brain plasticity. Our results show opposite and significant differences, consistent with their very different life-histories, in the sensory investments between these two closely-related species. Social parasites initially invest in the optic lobes to likely locate their hosts. After host colony usurpation, the parasite increases its brain volume, with specific investment in antennal lobes, which mirrors the behavioral switch from a usurping parasite to an integrated parasitic queen of the host colony. Contrastingly, hosts initially invest in the antennal lobes and sensory processing compared to social parasites, as predicted by their need to maintain social cohesion, allocate colony tasks, and recognize con- and heterospecific intruders. Host queens show a trend of higher investment in all sensory brain regions compared to workers, paralleling differences in task allocations. Our work provides novel insights into how intraspecific brain plasticity can facilitate the unique sensory adaptations needed to perform specific tasks by the host or to transition from searching to successful host exploitation by the social parasite.

## INTRODUCTION

Brood parasitism, in which a parasitic individual takes advantage of the parental care of a host, is a reproductive strategy that has evolved independently in diverse lineages, and is well-represented in birds and social insects (Antonson et al., 2020; Buschinger, 2009; Cini et al., 2019; Kilner & Langmore, 2011). The significant costs of parasitism to host species result in a coevolutionary arms race, where the parasite must locate and successfully exploit parental care by the host, while the host, in turn, must recognize and reject potential parasites (Feeney et al., 2012; Hauber et al., 2006; Lenoir et al., 2001). In particular, deceiving hosts is critical to obligate social parasites, a type of insect brood parasite that has lost the worker caste and solely depends on exploiting their social host for brood care (Rabeling, 2020). Although the adaptive behaviors of host and obligate social parasites have been studied extensively (Cervo, 2006; Lhomme & Hines, 2019; Loope et al., 2017; Nehring et al., 2015), the evolution and function of sensory systems to facilitate their behavioral interactions and arms-race remain poorly understood (Aidala et al., 2012; Stevens, 2013). Hosts have large sensory repertoires that facilitate general foraging decisions, social interactions, task allocation and the recognition of intruders. In contrast, the main selective pressure on obligate social parasites is to find and then deceive their hosts. Therefore, fine-tuned sensory systems are critical to mediate both the enemy-recognition by the host, and the successful deception and exploitation by the social parasite (Stoddard & Hauber, 2017).

However, developing brain tissue needed to process sensory stimuli is energetically expensive (Kotrschal et al., 2013; Niven & Laughlin, 2008; O’Donnell et al., 2011). Therefore, variation in the demands of specific sensory stimuli drives differential investment in specific sensory brain regions (Arganda et al., 2020; Barton et al., 1995; Keesey et al., 2020). For example, nocturnal lineages of mammals, birds, and insects have larger olfactory over visual brain structures compared to diurnal lineages (Barton et al., 1995; Corfield et al., 2008; O’Donnell et al., 2015; Sheehan et al., 2019; Stöckl et al., 2016). In social insects, specifically, differential investment in brain regions is also associated with colony size, caste, social interactions, and the need for distinguishing colony members from intruders (Arganda et al., 2020; Ehmer et al., 2001; O’Donnell et al., 2007; Seid et al., 2008; Seid & Junge, 2016).

Therefore, in systems where brood parasites attack social insects, hosts use their specialized sensory systems to recognize and reject intruders (Cervo et al., 1996; Cini, Bruschini, Signorotti, et al., 2011; Lenoir et al., 2001). In turn, parasites would need to use their own sensory systems to first locate potential hosts (Cervo et al., 1996; Cervo & Turillazzi, 1996), and then identify the correct host species, replace the host queen, and exploit the host workers for brood care (Cini, Bruschini, Poggi, et al., 2011; Nehring et al., 2015; Ortolani et al., 2010).

To explore adaptive investment in specific brain regions by hosts and obligate brood parasites, we here take advantage of a unique system composed of two closely-related paper wasp species (Choudhary et al., 1994). *Polistes sulcifer* is the obligate social parasite of *Polistes dominula* (Cervo & Turillazzi, 1996; Cini, Bruschini, Poggi, et al., 2011). Paper wasps use cuticular hydrocarbons (CHCs) as odor signals that indicate fertility and dominance (Dapporto et al., 2007), and to distinguish nestmates from intruders (Bruschini et al., 2011; Dani, 2006; Dani et al., 2001; Lorenzi et al., 1997; Mora-Kepfer, 2014). Therefore, precise and swift identification of potential parasites by CHCs recognition and visual inspection of facial markings is crucial to the host *P. dominula* (Cervo et al., 2015; Cini, Ortolani, et al., 2015; Ortolani et al., 2010; Sledge et al., 2001; Turillazzi et al., 2000). Paper wasp hosts queens and workers also have distinct sensory needs according to division of labor. Workers forage to collect nest material and prey, while queens remain on the nest and interact with these incoming subordinate workers (Queller et al., 2000).

In contrast, the social parasite *P. sulcifer* must overcome a different and complex challenge that requires a switch both in behavior and sensory modalities. A *P. sulcifer* female emerges from its overwintering site at the top of high mountains and migrates to lower elevations to locate the nests of *P. dominula* (Cervo, 2006), requiring navigational and visual acuity. After finding a host colony, the parasite usurps and functionally replaces the host queen to take reproductive control (Cervo, 2006), and acquires the CHCs of the colony (Bagnères & Lorenzi, 2010; Dapporto et al., 2004; Sledge et al., 2001; Turillazzi et al., 2000). However, host workers may eventually detect the new parasite queen, perhaps due to an imperfect chemical and/or behavioral integration into the host colony (Cini et al., 2020; Cini, Bruschini, Poggi, et al., 2011; Sledge et al., 2001). Nonetheless, when successful, the parasite queen becomes the sole egg layer (Cervo, 2006; Turillazzi et al., 2000). After the adult female and male parasite brood emerge from the host nest, they migrate to the top of mountains to mate, and fertilized females overwinter to start the search and usurpation cycle the following spring (Cervo, 2006). Therefore, *P. sulcifer’s* sensory needs should switch from an initial investment in vision needed to first migrate to mate and overwinter and migrate to locate a host colony the following spring, to a subsequent investment in olfaction to chemically deceive the host workers.

We hypothesize that relative proportions of select brain regions reflect the differential investment by hosts and parasites to meet their specific sensory needs. In particular, here we focus on insect brain regions with known functions. First, the antennal lobes (AL) receive olfactory stimuli from the antennae (Anton & Homberg, 1999). Second, the optic lobes (OL) are known to process visual input from the eyes and are associated to visual ecology needs (Gronenberg & Hölldobler, 1999). The OL are divided into the lamina (LA), medulla (ME) and lobula (LO), which provide contrast enhancement, color vision and motion detection, and shape discrimination, respectively (Arganda et al., 2020; Strausfeld, 1989; Yang et al., 2004; Yilmaz et al., 2019). Third, the mushroom bodies (MB) are the neuropils associated to learning and memory (Ehmer & Hoy, 2000; Strausfeld et al., 1998). In the calyx (CA) of the MB, the lip (LI) and collar (CO) primarily processes olfactory visual information, respectively (Ehmer & Hoy, 2000; Strausfeld et al., 1998). Finally, the Central complex (CX), the navigation center of the brain, may be implicated in long distance-migrations (Honkanen et al., 2019).

Given the known functions of these specific brain regions and the broad knowledge about the natural history of this social parasite-host system, we first predicted that *P. sulcifer* would show greater investment in the OL and CX compared to its host due to their need to migrate to overwinter and then find new hosts in the spring. Second, we explored if brain plasticity mirrors the drastic change from finding a host to integrating as a parasite queen. We predicted that parasites initially invest in brain structures necessary to navigate and locate a host and then switch to an investment in sensory structures after successfully integrating into a host nest. Third, we expected that hosts would show higher AL investment compared to its parasite, due to the different and complex olfactory stimuli they encounter and must assess (Dani, 2006). Finally, because social interactions increase the CA volume in *Polistes* wasps (Ehmer et al., 2001; Molina & O’Donnell, 2007), queen and worker hosts would show higher investment in the LI and CO, compared to the social parasite before it integrates into the host colony.

## METHODS

### Field collection and usurpation experiments

In Spring 2016 and 2017, we collected host colonies from unparasitized populations in the surroundings of Florence (Tuscany, Italy). Colonies had 2-4 foundresses and brood, but no adult workers. We fixed each nest to the ceiling of a glass cage (15 cm x 15 cm x 15 cm) and maintained it under controlled laboratory conditions with ad libitum sugar, water, fly maggots as larvae food, and paper for nest building. We individually marked colony members with enamel paint dots (Testor ©) on the thorax and wings. We also collected *P. sulcifer* females emerging from their overwintering sites at 2050 m altitude on Monti Sibillini (Umbria-Marche, Italy). We kept them in the same type of glass cages, inside a fridge at 4°C with ad libitum water and sugar, until it was time to end diapause. We simulated spring conditions by exposing the parasites to direct natural and artificial sunlight, and low-elevation natural temperature according to the established protocol (Cini et al., 2020; Cini, Bruschini, Poggi, et al., 2011; Ortolani et al., 2008)

We ran experimental trials during the last week of May, when usurpation usually occurs in the field (Cervo et al., 1996; Ortolani et al., 2008; Turillazzi et al., 1990). We focused on four experimental categories: usurping parasites, post-usurpation parasites, and host and workers in unparasitized colonies. For usurpation trials, we randomly chose host colonies and introduced a single social parasite female inside the glass cage of a putative host nest (Cini, Bruschini, Poggi, et al., 2011). We only chose parasites that showed usurpation behavior, confirmed by the clear attempts to land on the colony and the attacks toward the host foundresses. First, we confirmed that a parasite was trying to usurp a colony, and instead of allowing to integrate into the host colony, it was collected and categorized as a usurping parasite (N =4). Second, we allowed a subset of landing parasites to remain on the host nest. One week after usurpation, we confirmed that parasites showed typical behaviors of integrated parasite queens, such as occupying the central position of the nest, lack of foraging, stroking its abdomen on the nest cell rims, and dominating host individuals (Turillazzi et al., 1990). We categorized them as post-usurpation parasites (N =6). Finally, we reared unparasitized host colonies under the same conditions for one week after the emergence of workers. Each worker was marked with paint individually. We then collected the behaviorally dominant female and categorized it as host queen (N =7). From the same colony, we also collected a host worker, after confirming it was active and exhibiting typical worker behavior (N =4). To check that these behavioral categories matched the predicted reproductive physiology, we dissected the abdomens and assessed ovary development of each specimen using the established method for *Polistes* (Barth et al., 1975; Cini et al., 2013; Pardi, 1948; Walton et al., 2020). Therefore, we confirmed that host queens and parasite queens had developed ovaries, while host workers and usurping parasites had undeveloped ovaries.

### Histology and quantification of brain structures

We employed an established histological protocol for *Polistes* brains to test for differences in investment in brain structures that receive and process sensory information (Ehmer & Hoy, 2000; Molina & O’Donnell, 2008; O’Donnell et al., 2011; O’Donnell et al., 2007; O’Donnell et al., 2019). Each specimen was imbedded in epoxy resin to preserve the dimensions of the head capsule and the brain regions to avoid confounding measurements due to changes in brain size (Ocampo et al., 2020). To facilitate subsequent quantification of each brain region, heads were sectioned into 17 µm-thick coronal axis sections and stained with the NISSL stain toluidine blue. We then photographed each consecutive brain section per specimen by using a Canon camera (EOS 5D Mark III) mounted on a Leica DM IL LED microscope at 4x magnification and a 1000-micron scale. We used the image software AxioVision Software (version 4.8; Zeiss) to trace the AL, and the three substructures of the OL that receive visual information (LA, ME and LO). We also traced two CA substructures: LI and CO. Finally, we traced the CX, and the central brain (CB). We traced every other section in each specimen, as established by the reported accuracy of less than 3.5% error for 34 micro meter thick sections (Ehmer et al., 2001). Furthermore, we traced one hemisphere of the brain for each specimen, quantified area and calculated volume of each brain region (Molina & O’Donnell, 2008; Molina & O’Donnell, 2007; O’Donnell et al., 2011). Tracing and quantification of brain structures were performed blindly to the 2 species and 4 categories.

Each image was standardized using a 100 µm scale, and head width was measured as a proxy for body size. Finally, we used the software RECONSTRUCT to generate the 3D brain reconstructions for these two species (Fiala, 2005).

### Statistical analyses

Observed differences in proportional investment in different brain regions can arise from changes in allometric scaling through grade shifts (e.g, change in elevation) and/or changes in the slope of the covariance between brain regions (Eberhard & Wcislo, 2011; O’Donnell et al., 2013; Ott & Rogers, 2010; Seid et al., 2011; Sheehan et al., 2019; Stöckl et al., 2016). Thus, we here explored 1) if hosts and parasites differed in the relative size of specific brain regions, and 2) if changes in grade shift and/or slope explained the investment in specific sensory regions compared to non-sensory regions. To test for differences in visual and olfactory investment between host and social parasites, we first quantified absolute volume for each brain region and for the WB. For the specimens used in this study, the OL contribute on average to 42% of the total brain in *P. dominula* and 46% in *P. sulcifer*, which can influence the scaling of relative brain regions. Therefore, we normalized individual brain regions by CB volume to control for the effect of the OL in relative neuropil scaling, and avoid distortions of brain volume and size per species (Ott & Rogers, 2010; Sheehan et al., 2019; Stöckl et al., 2016).

To determine the relationship in the investment between sensory brain regions and central brain, we implemented Standardized Major (SMA) regression analyses, using the SMATR v.3 package for R (Warton et al., 2012; Warton et al., 2006). We utilized the scaling relationship between brain regions x and y, using the allometric equation y =a*x^β^ (Dubois, 1897; Huxley & Teissier, 1936). We then used the linear equation log(y) = βlog(x) + log(a), where log(a) = *α*, as this logarithmic transformation estimates β from the slope and *α* from intercept of a regression (Huxley & Teissier, 1936), used in previous studies that calculated investment in brain regions (Ott & Rogers, 2010; Sheehan et al., 2019; Stöckl et al., 2016).

First, we tested for a Common Slope between host and social parasite (H^0^ = *β*_*host*_ *= β*_*parasite*_) by using a log-likelihood test. Specifically, we ran the following comparisons in neural tissue investment: 1) WB and body size, 2) CB and WB, 3) pooled sensory regions and CB, and 4) each sensory region normalized by the CB. Second, if the host and parasite shared a Common Slope, we then tested for a Slope Index (SI), a Common Shift and a Grade Shift Index (GSI) for the four comparisons described above. The Slope Index (SI = *β*_host&par_) tested if investment in a brain region was isometric, calculated by a log-likelihood test. Therefore, if *β* −= 1, the proportion of sensory brain region (brain region y) and central brain (x) is allometric, meaning that x/y would change with size. The Common Shift (H^0^ = equal axis between host and parasite) indicated any shift along the major axis, calculated by a Wald Test. The Grade Shift Index (GSI) quantified how much larger a sensory brain region (region y) is for a given size of central brain (region x) for hosts compared to parasites (H^0^ = *α* _*host*_ *= α* _*parasite*_). GSI represents changes in elevation (intercept *α*) with no changes in the slope (*β*), reflecting volumetric differences between the two species as e ^*α host − α parasite*^, calculated by a Wald test. If the GSI > 1, hosts had a larger brain region than parasites, and if GSI< 1, parasites had a larger brain region than hosts. Log-likelihood tests and Wald tests were implemented in the SMATR package. Finally, we compared investment between the host and social parasite in specific sensory regions by normalizing each brain region by the CB, and running Mann-Whitney U tests. We ran these same comparisons among queen hosts, worker hosts, usurping parasites and parasite queens by using Kruskal-Wallis tests with posthoc pairwise comparisons, adjusted with Bonferroni corrections.

## RESULTS

### Contribution of allometry to host and social parasite differences

Hosts and parasites did not share a common allometric slope when we compared investment between absolute WB and body size (*P* =0.026, Fig. 1A, Suppl. Table 1). Hosts showed an isometric pattern with workers having small WB and body size while host queens showed a large WB and body size. In contrast, the social parasites showed a hyperallometric relationship in brain investment in the transition from usurpation to post-usurpation. Specifically, usurping parasites had small WB, but parasite queens increased their WB volume after successful integration into host colonies (Fig. 1A). Both the relationship between WB and CB (Fig 1. B), and between CB and pooled sensory brain regions (Fig 1.C) (See Suppl. Table 1) shared a common slope, respectively. Hosts showed a grade shift, with increased CB volume compared to the usurping social parasites, as indicated by significant differences in elevation (GSI >1, *P*=0.014). Parasites also showed higher investment in volume of pooled sensory brain regions compared to hosts, with the OL representing > 65% of these structures (GSI<1, *P*=0.016).

**Fig. 1.**
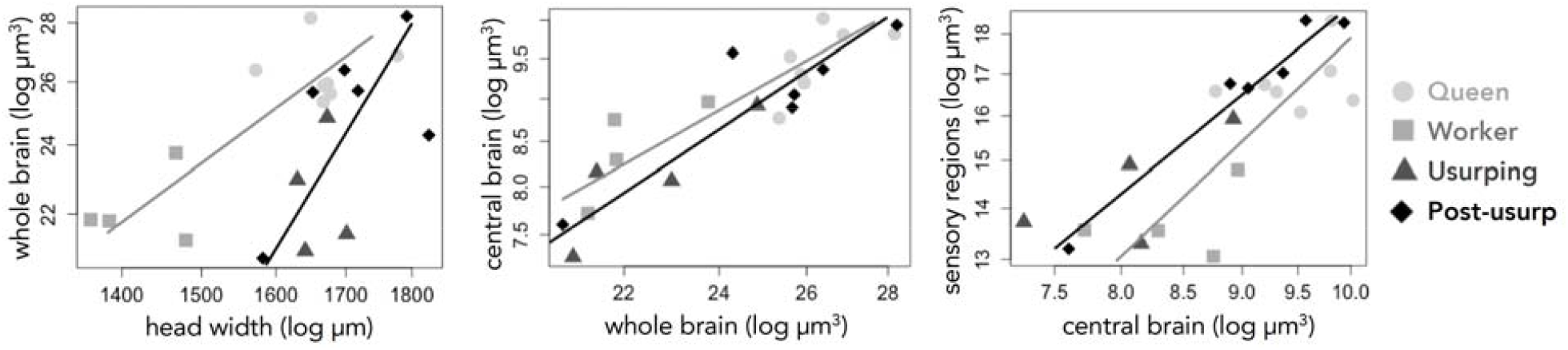
Role of allometry in investment of whole brain (WB), central brain (CB) and combined sensory brain regions between host and social parasite. Relationship between A) head width (as a proxy for body size) and WB, B) WB and CB and C) CB and combined sensory regions. Species and categories per species are depicted as: host queens (light grey circles) and workers (medium grey squares), usurping parasites (dark grey triangles) and post-usurpation parasite queens (black diamonds). Standardized major axis fits are log transformed per species with the lines based on intercepts and slopes (grey for hosts and black for social parasites). For full statistical tests, see Supplementary Table 1.

Notably, there is overlap in investment of pooled sensory regions between host queens and parasite queens (Fig. 1C). We found no effect of body size on brain region volume (SI) in the CB or pooled sensory regions (*P*=0.25 and *P*=0.9, respectively, Suppl. Table 1).

Next, hosts and social parasites shared a common slope in the relationship between each sensory brain region and CB, but showed unique differences in the GSI and SI depending on the specific region (Suppl. Table 1, Suppl. Figure 2). The LA that provides visual contrast enhancement, was significantly larger in parasites compared to hosts (GSI <1, *P*<0.001) and allometric, as indicated by significant differences in the SI (*P*<0.001). The ME and LO, that facilitate color vision, motion detection and shape discrimination, also had increased volume in parasites compared to hosts (GSI <1, *P*<0.001 and *P*=0.01, respectively), but were isometric.

However, the AL showed an opposite and allometric increase in the hosts compared to the parasites. Both the CO and LI, that process visual and olfactory stimuli respectively, also showed this allometric pattern. The LI also had a significant grade shift (GSI>1, *P* <0.001) in the hosts compared to parasites. Finally, the CX, the navigation center of the brain, had significantly increased volume in parasites compared to hosts (GSI <1, *P*=0.045).

### Differential relative investment and tradeoffs between sensory brain regions

When comparing the relative investment in sensory regions normalized by the CB, we found significant and opposite differences between hosts and social parasites. Hosts had increased AL volume (U=19.00, df =1, *P*=0.011, Fig 2A,) and a larger CA than social parasites (U=17.00, df =1, P=0.007, Suppl Fig. 1) Both CA substructures, LI and CO, were significantly larger in the host than in the parasite (U=8.00, df =1, P=0.001 and U=9.76, df=1, P=0.03 respectively, Fig. 2B). In contrast, OL volume was larger in social parasites compared to hosts (U=103.00, df =1, *P*=0.001, Suppl. Fig 1A). Specifically, social parasites invested highly in the LA (U=105.00, df =1, P<0.001, Fig. 2D), ME (U=95.00, df =1, P=0.005, Fig. 2D) and LO (U=91.00, df =1, P=0.011, Fig. 2D). Social parasites also showed higher volume in the CX, compared to the hosts (U=79.00, df =1, P=0.025, Fig. 2E).

**Figure 2.**
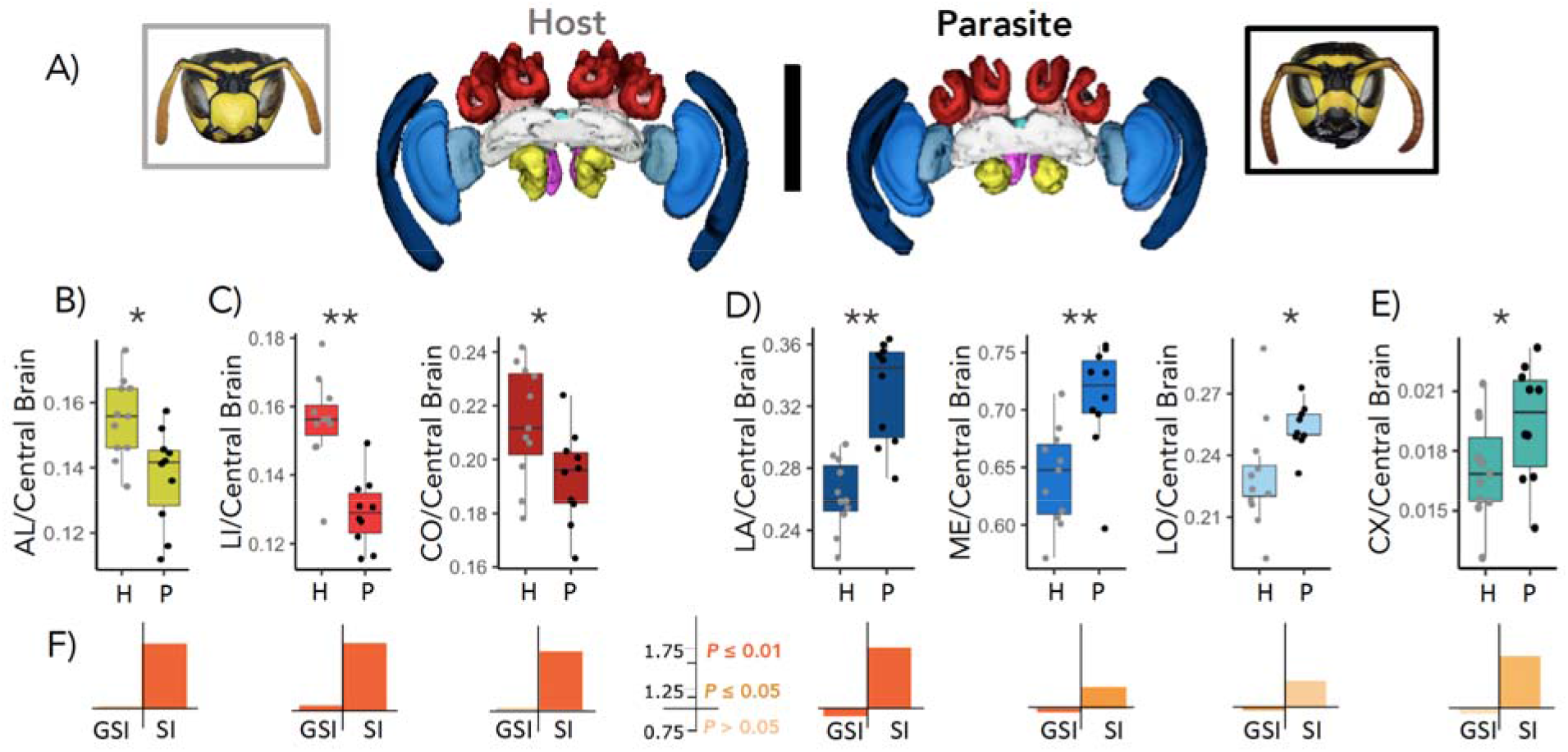
Visual and olfactory brain regions in *Polistes dominula* (host) and *P. sulcifer* (social parasite). A) Frontal view of 3D reconstructed brain regions for the host (grey and to the left) and the parasite (black and to the right). Black scale bar = 1 mm. B) Antennal lobes (AL). C) Substructures of the calyx (CA): lip (LI) and collar (Co). D) Substructures of the OL: lamina (LA), medulla (ME) and lobula (LO) in sequence. E) Central complex (CX). All brain regions are normalized by the central brain shown in light grey. The subesophageal zone (ZEZ) is shown in dark pink and the peduncle (PED) in light pink. Each dot represents an individual and is grey for hosts (H) and black for parasites (P). Each box plot shows the median, 25^th^ and 75^th^ percentiles and the whiskers show the 5^th^ and 95^th^ percentiles. E) The Grade Shift Index (GSI) was calculated by scaling differences in normalized sensory brain regions between species. If GSI > 1, hosts had higher investment than parasites, and vice versa for GSI < 1. The Slope Index (SI) is represented by the deviation of the estimated common allometric slope *β* from 1. Statistical results based on Mann-Whitney U tests (^*^ *P* < 0.05, ^**^ *P* < 0.01). Full statistical tests can be found in the Results sections and Supplementary Table 1. See Suppl. Fig 1. for OL and CA results.

We then tested for differential investment in sensory brain regions among the four categories: 1) host queens and 2) host workers from the same nest, 3) usurping parasites, 4) post-usurpation (parasite queens). Both usurping parasites and parasite queens had significantly larger OL volume, followed by the host queen, while the host workers had the smallest OL (χ2=12.36, df =3, P =0.006, Suppl. Fig. 1). We found the same pattern for investment among categories in the LA (χ^2^=12.79, df =3, P =0.005, Fig. 3), ME (χ ^2^=8.89, df =3, P =0.031) and LO χ =10.14, df =3, P =0.017, respectively, Fig. 3C). Contrastingly, host queen had the highest investment in AL while usurping parasites had the lowest investment. However, parasite queens had a similar AL volume to host workers, which reflects an increase in investment after usurpation (χ^2^=11.11, df =3, P =0.01, Fig 3A). Usurping parasites and parasite queens had significantly smaller CA compared to host queens. However, parasite queens had similar CA volume to host workers, showing a trend towards increased investment in this brain region after usurpation (χ^2^=12.14, df =3, P =0.007). Host queens also had a significantly larger LI volume,followed by an intermediate volume in host workers, and small LI in usurping parasites and parasite queens. (χ^2^=13.12, df =3, P =0.004, Fig. 3B). Host queens and workers had the largest CO, while usurping parasites had relatively small CO. However, parasite queens showed a significant increase in CO volume, compared to (χ^2^=9.76, df =3, P =0.021, Fig. 3B).

**Figure 3.**
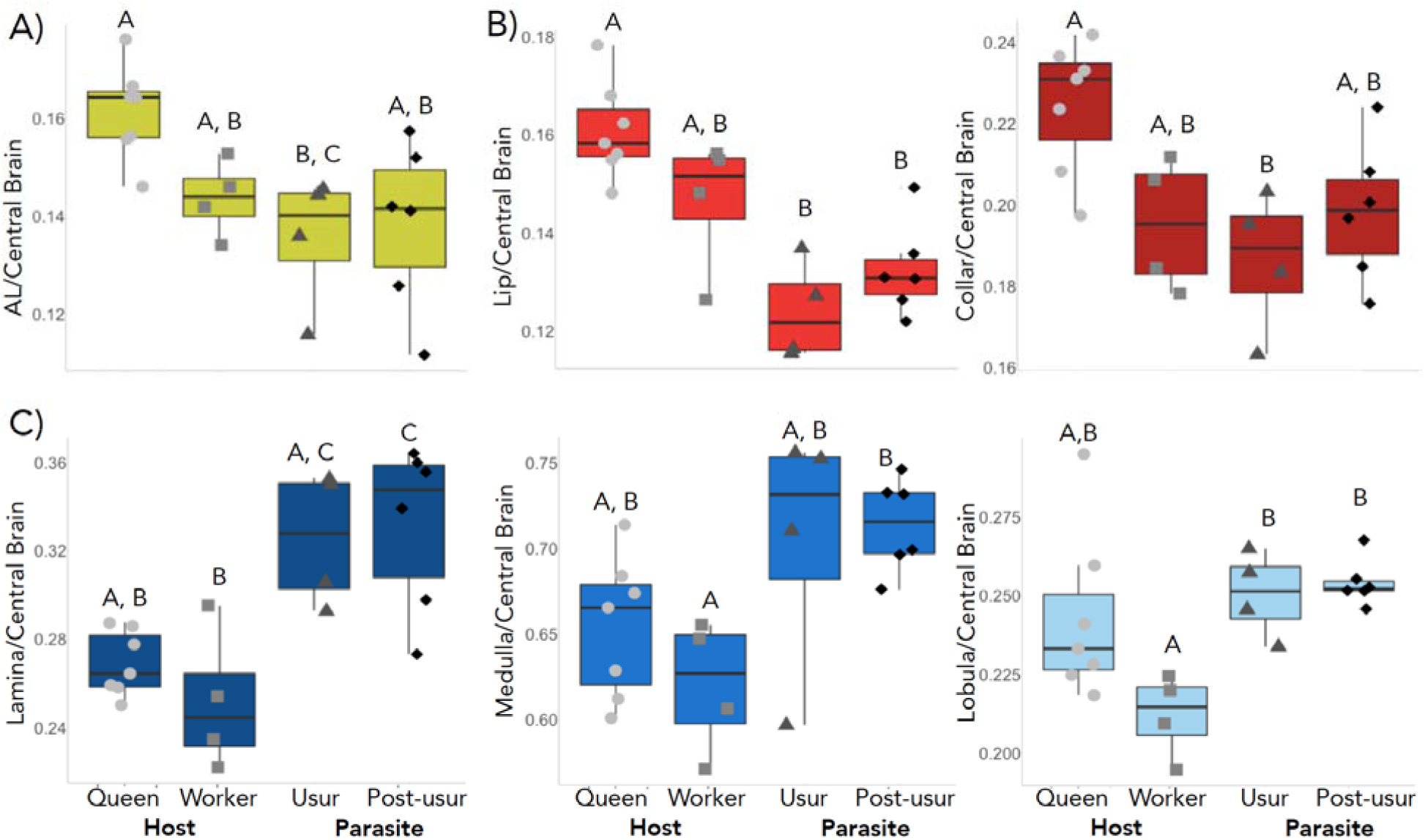
Visual and olfactory brain regions in host queens (light grey circles) and workers (medium grey squares) and usurping parasites (dark grey triangles) and post-usurpation parasite queens (black diamonds). A) Antennal lobes (AL), B) Substructures of the calyx (CA): lip (LI) and collar (CO), C) Substructures of the optic lobes (OL): lamina (LA), medulla (ME) and lobula (LO) in order from dark to light blue. Each dot represents an individual and each box plot shows the median, 25^th^ and 75^th^ percentiles and the whiskers show the 5^th^ and 95^th^ percentiles. Results reported by Mann-Whitney U tests, with different letters stating significant differences between categories. Full statistical tests can be found in the results section.

## DISCUSSION

Social parasites have evolved sensory and behavioral strategies to effectively locate host nests and integrate as the new parasitic queen, while wasp hosts have evolved complex behavioral repertoires needed to recognize nestmates and intruders (Bagnères & Lorenzi, 2010; Brandt et al., 2005; Cervo, 2006; Cini, Patalano, et al., 2015; Cini et al., 2019). In this study, hosts and parasites showed an inverse relationship in the investment towards visual and olfactory brain regions, which supports the hypothesis that expensive brain tissue is allocated according to specific sensory needs (Keesey et al., 2020). For instance, the most important selective pressure in the social parasite is to first find a host nest, then functionally replace the host queen and deceive the host workers (Cini, Patalano, et al., 2015; Cini et al., 2019; Grüter et al., 2018), which requires complex changes in sensory processing. We find several lines of evidence supporting the hypothesis that brain plasticity facilitates these crucial behavioral and sensory transitions in the social parasite. First, *P. sulcifer* shows high investment in the optic lobes, which coincides with the selective and energetic pressures for migration, host detection and invasion, as the visual system is expensive to maintain (Niven & Laughlin, 2008). Second, during the usurpation period, there is less investment in the antennal lobes and both visual and olfactory processing by the calyx. Third, our results suggest that brain plasticity is associated with the switch from a searching solitary parasite to invading a host colony and disguising itself as a parasite queen. The social parasite significantly increases whole brain volume as it transitions from usurping to post-usurpation, reflected in the significant increase in antennal lobes and a trend in increase of the calyx. In sum, our results suggest that brain plasticity is advantageous, allowing parasites to adaptively respond to changes in sensory needs (Beani et al., 2014; Molina & O’Donnell, 2008; Molina & O’Donnell, 2007).

Notably, energetic allocation towards the optic lobes mirrors the initial visual needs of this obligate social parasite. Investment in the lamina, medulla and lobula may enhance contrast, motion detection, color vision and discrimination (Arganda et al., 2020; Gronenberg et al., 2008; Strausfeld, 1989; Yang et al., 2004; Yilmaz et al., 2019), which may be critical during migration to first overwinter in its mountain site, and then migrate again to find suitable host nests to invade at lower elevations (Cervo, 2006). In addition, the parasite uses visual cues to localize nests, and then olfactory cues to distinguish *P. dominula* from sympatric *P. gallicus* or *P. nimpha*, as the parasite’s large size can hinder effective usurpation in the latter two social species (Cervo, 2006; Cervo & Turillazzi, 1996; Cini, Bruschini, Signorotti, et al., 2011). The high investment in vision and low investment in sensory processing (e.g., calyx) also reflects the initial needs of the parasite, which is remarkably similar to neural tissue allocation in the obligate parasitic ant *Polyergus mexicanus* (Sulger et al., 2014). In addition, the central complex, the navigation center of the brain, is significantly larger in the social parasite compared to the host (Honkanen et al., 2019).

After successfully finding a host nest then replacing the host queen, the parasite experiences a drastic change in sensory needs from its usurping phase. We find that the change is reflected by an increase in whole brain volume, which is explained by reallocation of investment in the antennal lobe and a trend towards increasing sensory processing. This pattern suggests the need to adopt the CHCs of the colony to become reproductively dominant (Turillazzi et al., 2000). If the parasite succeeds, host workers will treat the parasite as if it were the host queen (Cervo, 2006; Cini, Patalano, et al., 2015). However, recent experimental works suggests that host workers eventually recognize the parasite queen as an intruder, so the time to produce and rear successful parasite offspring is limited (Cini et al., 2020; Cini et al., 2014). Behavioral and neurogenomic responses by host workers (Cini et al., 2020) may elicit the investment by the parasite queen to increase the antennal lobes to receive olfactory information, and both in the lip and collar, which process olfactory and visual information respectively. Once the parasite integrates itself as parasite queen, this increase in the calyx is similar to the investment by host workers, which is associated to social interactions in *Polistes* wasps (Ehmer et al., 2001; Molina & O’Donnell, 2008; Molina & O’Donnell, 2007). However, calyx investment in parasite queens is less than in host queens, which may be explained by differences in their behavioral repertoires (Cini, Patalano, et al., 2015), with host queens having more frequent social interactions.

In contrast, parasite attacks do not represent the strongest selective pressure in the sensory system of hosts, as incidence of obligate parasitism is almost null in this host population (RC & AC, pers. comm.). Our results of high investment in olfaction and sensory processing are consistent with other studies that show preferential brain investment in brain region associated needed for effective communication, maintenance of division of labor, learning and memory (Farris et al., 2001; Gronenberg et al., 1996; Jaumann et al., 2019; O’Donnell et al., 2015; Rehan et al., 2015; Seid et al., 2005; Smith et al., 2010). Host queens had significantly higher antennal lobes, as they spend most of their time on the nest and communicate with incoming subordinate workers, who spend more time foraging off the nest (Molina & O’Donnell, 2008). Host queens also use olfaction to assess fertility and policing attempts by subordinate workers, and to maintain their dominance in the nest hierarchy (Dapporto et al., 2010). As predicted, host queens also had more developed antennal lobes than usurping parasites and parasite queens. The antennal lobes in *P. dominula* may facilitate identifying non-nestmate conspecifics or social parasites, as recent work in paper wasps shows both an expansion and rapid evolution in the 9-exon odorant receptors, responsible for detecting CHCs (Legan et al., 2020). Because *P. sulcifer* must adopt the odor of the host colony to become chemically integrated, it is critical for the host queen to detect any intruders approaching the nest, including social parasites (Cervo, 2006; Cini et al., 2020; Turillazzi et al., 2000). Consistent with the critical task of recognition, we found that both host queens and workers invested more in brain regions for olfactory processing compared to obligate parasites. Hosts also invested more in visual processing compared to parasites, as indicated by higher collar volume. These findings suggest that differential brain investment may be a response to the important role of olfaction and vision in discriminating nest intruders, which has been shown experimentally in this host-parasite system (Cini, Ortolani, et al., 2015).

Another possibility is that high investment in the calyx by hosts is due to the visual and olfactory demands of foraging. Established parasitic queens, in contrast, will not forage and instead remain on the host nest (Cervo, 2006). This pattern is consistent with the observed higher calyx investment for the host ant *Formica fusta* compared to workers of its obligate social parasite *P. mexicanus*, which rarely forage (Sulger et al., 2014). These two hypotheses may not be mutually exclusive, and further experiments that include molecular, cellular and circuitry approaches (Godfrey & Gronenberg, 2019) would elucidate the effect of these two selective pressures towards sensory investment.

## Conclusions

Our study provides novel insights into how intraspecific brain plasticity facilitates the preferential investment of specific neural structures to meet an individual’s sensory needs. In the obligate parasite, our results support the hypothesis that preferential investment in specific brain regions reflects the changing sensory needs of an individual, as it switches from searching for a host colony to invading and disguising itself as a parasite queen. Specifically, the social parasite may first invest in the optic lobes to facilitate migration and locating host nests, and then transition to an increase in the antennal lobes to successfully integrate into the host colony. Given that this social parasite evolved from a eusocial ancestor, it is likely retaining traits that facilitate this cognitive switch from a solitary parasite to a social parasite queen in the host colony. Previous work suggest that social insects share genetic toolkit genes (Berens et al., 2014; Toth et al., 2010), so the social parasites likely respond to changes in sensory needs similarly to their eusocial relatives (Cini, Patalano, et al., 2015; Cini et al., 2019). In the potential host, our data suggests that queens and workers take advantage of their existing, and well-developed sensory system, and particularly the antennal lobes, for intruder detection to increase social parasite rejection. A trend towards differential investment in sensory structures is also mirrored in the differential task allocation between host queens and workers. Finally, future work comparing host populations highly attacked or not attacked by *P. sulcifer* will effectively test the role of these social parasites as a selective pressure on the differential investment in specific brain sensory regions in their hosts.

## Supporting information

Rozanski_etal_Suppl_Material_NeuroAnSocPar

## ACKNOWLEDGEMENTS

Sean O’Donnell, William Searcy, Chris Jernigan, Al Uy and members of the Uy labs provided thoughtful comments in previous versions of this manuscript. We thank Irene Pepiciello and Federico Cappa for help in maintaining the wasps in laboratory conditions and performing usurpation experiments, Diego Ocampo for statistical advice and Marco Selis for providing the host and parasite photographs. ANR was supported by CAS Summer Research Program for Underrepresented Minorities & Women at the University of Miami. ANR, FMKU and MEH were supported by a National Academies Keck Future Initiatives Grant (NAKFI). RC and AC were supported by University of Florence funds. Samples from Italy were imported into the USA via USDA permit #128388.

## LITERATURE CITED

Aidala, Z., Huynen, L., Brennan, P. L., Musser, J., Fidler, A., Chong, N., Machovsky Capuska, G. E., Anderson, M. G., Talaba, A., Lambert, D., & Hauber, M. E. (2012). Ultraviolet visual sensitivity in three avian lineages: paleognaths, parrots, and passerines. Journal of Comparative Physiology A, 198(7), 495–510.

Anton, S., & Homberg, U. (1999). Antennal lobe structure. In Insect Olfaction (pp. 97–124). Berlin, Heidelberg: Springer.

Antonson, N. D., Rubenstein, D. R., Hauber, M. E., & Botero, C. A. (2020). Ecological uncertainty favours the diversification of host use in avian brood parasites. Nature Communications, 11(1), 1–7.

Arganda, S., Hoadley, A. P., Razdan, E. S., Muratore, I. B., & Traniello, J. F. A. (2020). The neuroplasticity of division of labor: worker polymorphism, compound eye structure and brain organization in the leafcutter ant Atta cephalotes. Journal of Comparative Physiology A, 206(4), 651–662.

Bagnères, A.-G., & Lorenzi, M. C. (2010). Chemical deception/mimicry using cuticular hydrocarbons. In Insect hydrocarbons: biology, biochemistry and chemical ecology (pp. 282–324). Cambridge: Cambridge University Press.

Barth, R., Lester, L., Sroka, P., Kessler, T., & Hearn, R. (1975). Juvenile hormone promotes dominance behavior and ovarian development in social wasps (Polistes annularis). Experientia, 31(6), 691–692.

Barton, R., Purvis, A., & Harvey, P. (1995). Evolutionary radiation of visual and olfactory brain systems in primates, bats and insectivores. Philosophical Transactions of the Royal Society of London. Series B: Biological Sciences, 348(1326), 381–392.

Beani, L., Dessì-Fulgheri, F., Cappa, F., & Toth, A. (2014). The trap of sex in social insects: from the female to the male perspective. Neuroscience & Biobehavioral Reviews, 46, 519–533.

Berens, A. J., Hunt, J. H., & Toth, A. L. (2014). Comparative transcriptomics of convergent evolution: different genes but conserved pathways underlie caste phenotypes across lineages of eusocial insects. Molecular Biology and Evolution, 32(3), 690–703.

Brandt, M., Foitzik, S., Fischer-Blass, B., & Heinze, J. (2005). The coevolutionary dynamics of obligate ant social parasite systems–between prudence and antagonism. Biological Reviews, 80(2), 251–267.

Bruschini, C., Cervo, R., Cini, A., Pieraccini, G., Pontieri, L., Signorotti, L., & Turillazzi, S. (2011). Cuticular hydrocarbons rather than peptides are responsible for nestmate recognition in Polistes dominulus. Chemical Senses, 36(8), 715–723.

Buschinger, A. (2009). Social parasitism among ants: a review (Hymenoptera: Formicidae). Myrmecological News, 12(3), 219–235.

Cervo, R. (2006). Polistes wasps and their social parasites: an overview. Annales Zoologici Fennici, 531–549.

Cervo, R., Bertocci, F., & Turillazzi, S. (1996). Olfactory cues in host nest detection by the social parasite Polistes sulcifer (Hymenoptera, Vespidae). Behavioural Processes, 36(3), 213–218.

Cervo, R., Cini, A., & Turillazzi, S. (2015). Visual recognition in social wasps. In Social recognition in invertebrates (pp. 125–145). Cham (Switzerland): Springer International Publishing.

Cervo, R., & Turillazzi, S. (1996). Host nest preference and nest choice in the cuckoo paper wasp Polistes sulcifer (Hymenoptera: Vespidae). Journal of Insect Behavior, 9(2), 297–306.

Choudhary, M., Strassmann, J. E., Queller, D. C., Turillazzi, S., & Cervo, R. (1994). Social parasites in polistine wasps are monophyletic: implications for sympatric speciation. Proceedings of the Royal Society of London. Series B: Biological Sciences, 257(1348), 31–35.

Cini, A., Branconi, R., Patalano, S., Cervo, R., & Sumner, S. (2020). Behavioural and neurogenomic responses of host workers to social parasite invasion in a social insect. Insectes Sociaux, 67(2), 295–308.

Cini, A., Bruschini, C., Poggi, L., & Cervo, R. (2011). Fight or fool? Physical strength, instead of sensory deception, matters in host nest invasion by a wasp social parasite. Animal Behaviour, 81(6), 1139–1145.

Cini, A., Bruschini, C., Signorotti, L., Pontieri, L., Turillazzi, S., & Cervo, R. (2011). The chemical basis of host nest detection and chemical integration in a cuckoo paper wasp. Journal of Experimental Biology, 214(21), 3698–3703.

Cini, A., Meconcelli, S., & Cervo, R. (2013). Ovarian indexes as indicators of reproductive investment and egg-laying activity in social insects: a comparison among methods. Insectes Sociaux, 60(3), 393–402.

Cini, A., Nieri, R., Dapporto, L., Monnin, T., & Cervo, R. (2014). Almost royal: incomplete suppression of host worker ovarian development by a social parasite wasp. Behavioral Ecology and Sociobiology, 68(3), 467–475.

Cini, A., Ortolani, I., Zechini, L., & Cervo, R. (2015). Facial markings in the social cuckoo wasp Polistes sulcifer: No support for the visual deception and the assessment hypotheses. Behavioural Processes, 111, 19–24.

Cini, A., Patalano, S., Segonds-Pichon, A., Busby, G. B., Cervo, R., & Sumner, S. (2015). Social parasitism and the molecular basis of phenotypic evolution. Frontiers in Genetics, 6, 32.

Cini, A., Sumner, S., & Cervo, R. (2019). Inquiline social parasites as tools to unlock the secrets of insect sociality. Philosophical Transactions of the Royal Society B, 374(1769), 20180193.

Corfield, J. R., Wild, J. M., Hauber, M. E., Parsons, S., & Kubke, M. F. (2008). Evolution of brain size in the Palaeognath lineage, with an emphasis on New Zealand ratites. Brain, Behavior and Evolution, 71(2), 87–99.

Dani, F. R. (2006). Cuticular lipids as semiochemicals in paper wasps and other social insects. Annales Zoologici Fennici, 500-514.

Dani, F. R., Jones, G. R., Destri, S., Spencer, S. H., & Turillazzi, S. (2001). Deciphering the recognition signature within the cuticular chemical profile of paper wasps. Animal Behaviour, 62(1), 165–171.

Dapporto, L., Bruschini, C., Cervo, R., Dani, F. R., Jackson, D. E., & Turillazzi, S. (2010). Timing matters when assessing dominance and chemical signatures in the paper wasp Polistes dominulus. Behavioral Ecology and Sociobiology, 64(8), 1363–1365.

Dapporto, L., Cervo, R., Sledge, M., & Turillazzi, S. (2004). Rank integration in dominance hierarchies of host colonies by the paper wasp social parasite Polistes sulcifer (Hymenoptera, Vespidae). Journal of Insect Physiology, 50(2-3), 217–223.

Dapporto, L., Dani, F. R., & Turillazzi, S. (2007). Social dominance molds cuticular and egg chemical blends in a paper wasp. Current Biology, 17(13), R504–R505.

Dubois, E. (1897). Sur le rapport du poids de l’encéphale avec la grandeur du corps chez les mammifères. Bulletins et Mémoires de la Société d’Anthropologie de Paris, 8(1), 337–376.

Eberhard, W. G., & Wcislo, W. T. (2011). Grade changes in brain–body allometry: morphological and behavioural correlates of brain size in miniature spiders, insects and other invertebrates. In Advances in Insect Physiology (Vol. 40, pp. 155–214): Academic Press.

Ehmer, B., & Hoy, R. (2000). Mushroom bodies of vespid wasps. Journal of Comparative Neurology, 416(1), 93–100.

Ehmer, B., Reeve, H. K., & Hoy, R. R. (2001). Comparison of brain volumes between single and multiple foundresses in the paper wasp Polistes dominulus. Brain, Behavior and Evolution, 57(3), 161–168. doi:10.1159/000047234

Farris, S. M., Robinson, G. E., & Fahrbach, S. E. (2001). Experience- and age-related outgrowth of intrinsic neurons in the mushroom bodies of the adult worker honeybee. Journal of Neuroscience, 21(16), 6395–6404.

Feeney, W. E., Welbergen, J. A., & Langmore, N. E. (2012). The frontline of avian brood parasite–host coevolution. Animal Behaviour, 84(1), 3–12.

Fiala, J. C. (2005). Reconstruct: a free editor for serial section microscopy. Journal of Microscopy, 218(1), 52–61.

Godfrey, R. K., & Gronenberg, W. (2019). Brain evolution in social insects: advocating for the comparative approach. Journal of Comparative Physiology A, 205(1), 13–32.

Gronenberg, W., Ash, L. E., & Tibbetts, E. A. (2008). Correlation between facial pattern recognition and brain composition in paper wasps. Brain, Behavior and Evolution, 71(1), 1–14.

Gronenberg, W., Heeren, S. Ouml, & Lldobler, B. (1996). Age-dependent and task-related morphological changes in the brain and the mushroom bodies of the ant Camponotus floridanus. Journal of Experimental Biology, 199(Pt 9), 2011–2019.

Gronenberg, W., & Hölldobler, B. (1999). Morphologic representation of visual and antennal information in the ant brain. Journal of Comparative Neurology, 412(2), 229–240.

Grüter, C., Jongepier, E., & Foitzik, S. (2018). Insect societies fight back: the evolution of defensive traits against social parasites. Philosophical Transactions of the Royal Society B: Biological Sciences, 373(1751), 20170200.

Hauber, M. E., Moskát, C., & Bán, M. (2006). Experimental shift in hosts’ acceptance threshold of inaccurate-mimic brood parasite eggs. Biology Letters, 2(2), 177–180.

Honkanen, A., Adden, A., da Silva Freitas, J., & Heinze, S. (2019). The insect central complex and the neural basis of navigational strategies. Journal of Experimental Biology, 222. doi:jeb188854

Huxley, J. S., & Teissier, G. (1936). Terminology of relative growth. Nature, 137(3471), 780–781.

Jaumann, S., Seid, M. A., Simons, M., & Smith, A. R. (2019). Queen dominance may reduce worker mushroom body size in a social bee. Developmental Neurobiology, 79(6), 596–607.

Keesey, I. W., Grabe, V., Knaden, M., & Hansson, B. S. (2020). Divergent sensory investment mirrors potential speciation via niche partitioning across Drosophila. eLife, 9, e57008. doi:10.7554/eLife.57008

Kilner, R. M., & Langmore, N. E. (2011). Cuckoos versus hosts in insects and birds: adaptations, counter-adaptations and outcomes. Biological Reviews, 86(4), 836–852.

Kotrschal, A., Rogell, B., Bundsen, A., Svensson, B., Zajitschek, S., Brännström, I., Immler, S., Maklakov, A.A., & Kolm, N. (2013). Artificial selection on relative brain size in the guppy reveals costs and benefits of evolving a larger brain. Current Biology, 23(2), 168–171.

Legan, A. W., Jernigan, C. M., Miller, S. E., Fuchs, M. F., & Sheehan, M. J. (2020). Expansion and accelerated evolution of 9-exon odorant receptors in Polistes paper wasps. bioRxiv, 2020.2009.2004.283903. doi:10.1101/2020.09.04.283903

Lenoir, A., d’Ettorre, P., Errard, C., & Hefetz, A. (2001). Chemical ecology and social parasitism in ants. Annual Review of Entomology, 46(1), 573–599.

Lhomme, P., & Hines, H. M. (2019). Ecology and evolution of cuckoo bumble bees. Annals of the Entomological Society of America, 112(3), 122–140.

Loope, K. J., Lopez-Osorio, F., & Dvořák, L. (2017). Convergent reversion to single mating in a wasp social parasite. The American Naturalist, 189(6), E138–E151.

Lorenzi, M. C., Bagnères, A., Clément, J.-L., & Turillazzi, S. (1997). Polistes biglumis bimaculatus epicuticular hydrocarbons and nestmate recognition (Hymenoptera, Vespidae). Insectes Sociaux, 44(2), 123–138.

Molina, Y., & O’Donnell, S. (2008). Age, sex, and dominance-related mushroom body plasticity in the paperwasp Mischocyttarus mastigophorus. Developmental Neurobiology, 68(7), 950–959.

Molina, Y., & O’Donnell, S. (2007). Mushroom body volume is related to social aggression and ovary development in the paperwasp Polistes instabilis. Brain, Behavior and Evolution, 70(2), 137–144.

Mora-Kepfer, F. (2014). Context-dependent acceptance of non-nestmates in a primitively eusocial insect. Behavioral Ecology and Sociobiology, 68(3), 363–371.

Nehring, V., Dani, F. R., Turillazzi, S., Boomsma, J. J., & d’Ettorre, P. (2015). Integration strategies of a leaf-cutting ant social parasite. Animal Behaviour, 108, 55–65.

Niven, J. E., & Laughlin, S. B. (2008). Energy limitation as a selective pressure on the evolution of sensory systems. Journal of Experimental Biology, 211(11), 1792–1804.

O’Donnell, S., Clifford, M., & Molina, Y. (2011). Comparative analysis of constraints and caste differences in brain investment among social paper wasps. Proceedings of the National Academy of Sciences, 108(17), 7107–7112.

O’Donnell, S., Donlan, N., & Jones, T. (2007). Developmental and dominance-associated differences in mushroom body structure in the paper wasp Mischocyttarus mastigophorus. Developmental Neurobiology, 67(1), 39–46.

O’Donnell, S., Bulova, S., DeLeon, S., Barrett, M., & Fiocca, K. (2019). Brain structure differences between solitary and social wasp species are independent of body size allometry. Journal of Comparative Physiology A, 205(6), 911–916.

O’Donnell, S., Bulova, S. J., DeLeon, S., Khodak, P., Miller, S., & Sulger, E. (2015). Distributed cognition and social brains: reductions in mushroom body investment accompanied the origins of sociality in wasps (Hymenoptera: Vespidae). Proceedings of the Royal Society B: Biological Sciences, 282(1810), 20150791.

O’Donnell, S., Clifford, M. R., DeLeon, S., Papa, C., Zahedi, N., & Bulova, S. J. (2013). Brain size and visual environment predict species differences in paper wasp sensory processing brain regions (Hymenoptera: Vespidae, Polistinae). Brain, Behavior and Evolution, 82(3), 177–184.

Ocampo, D., Sánchez, C., & Barrantes, G. (2020). Do different methods yield equivalent estimations of brain size in birds? Brain, Behavior and Evolution, 95(2), 113–122.

Ortolani, I., Turillazzi, S., & Cervo, R. (2008). Spring usurpation restlessness: a wasp social parasite adapts its seasonal activity to the host cycle. Ethology, 114(8), 782–788.

Ortolani, I., Zechini, L., Turillazzi, S., & Cervo, R. (2010). Recognition of a paper wasp social parasite by its host: evidence for a visual signal reducing host aggressiveness. Animal Behaviour, 80(4), 683–688.

Ott, S. R., & Rogers, S. M. (2010). Gregarious desert locusts have substantially larger brains with altered proportions compared with the solitarious phase. Proceedings of the Royal Society B: Biological Sciences, 277(1697), 3087–3096.

Pardi, L. (1948). Dominance order in Polistes wasps. Physiological Zoology, 21(1), 1–13.

Queller, D. C., Zacchi, F., Cervo, R., Turillazzi, S., Henshaw, M. T., Santorelli, L. A., & Strassmann, J. E. (2000). Unrelated helpers in a social insect. Nature, 405(6788), 784–787.

Rabeling, C. (2020). Social Parasitism. In C. K. Starr (Ed.), Encyclopedia of Social Insects (pp. 1-23). Cham: Springer International Publishing.

Rehan, S. M., Bulova, S. J., & O’Donnell, S. (2015). Cumulative effects of foraging behavior and social dominance on brain development in a facultatively social bee (Ceratina australensis). Brain Behavior and Evolution, 85(2), 117–124.

Seid, M. A., Castillo, A., & Wcislo, W. T. (2011). The allometry of brain miniaturization in ants. Brain, Behavior and Evolution, 77(1), 5–13.

Seid, M. A., Goode, K., Li, C., & Traniello, J. F. (2008). Age-and subcaste-related patterns of serotonergic immunoreactivity in the optic lobes of the ant Pheidole dentata. Developmental Neurobiology, 68(11), 1325–1333.

Seid, M. A., Harris, K. M., & Traniello, J. F. A. (2005). Age-related changes in the number and structure of synapses in the lip region of the mushroom bodies in the ant Pheidole dentata. Journal of Comparative Neurology, 488(3), 269–277.

Seid, M. A., & Junge, E. (2016). Social isolation and brain development in the ant Camponotus floridanus. The Science of Nature, 103(5-6), 1–6.

Sheehan, Z. B., Kamhi, J. F., Seid, M. A., & Narendra, A. (2019). Differential investment in brain regions for a diurnal and nocturnal lifestyle in Australian Myrmecia ants. Journal of Comparative Neurology, 527(7), 1261–1277.

Sledge, M. F., Dani, F. R., Cervo, R., Dapporto, L., & Turillazzi, S. (2001). Recognition of social parasites as nest-mates: adoption of colony-specific host cuticular odours by the paper wasp parasite Polistes sulcifer. Proceedings of the Royal Society of London. Series B: Biological Sciences, 268(1482), 2253–2260.

Smith, A. R., Seid, M. A., Jiménez, L. C., & Wcislo, W. T. (2010). Socially induced brain development in a facultatively eusocial sweat bee Megalopta genalis (Halictidae). Proceedings of the Royal Society B: Biological Sciences, 277(1691), 2157–2163.

Stevens, M. (2013). Arms Race, Coevolution, and Diversification. In Sensory ecology, behaviour, and evolution (pp. 143–163). United Kingdom: Oxford University Press.

Stöckl, A., Heinze, S., Charalabidis, A., El Jundi, B., Warrant, E., & Kelber, A. (2016). Differential investment in visual and olfactory brain areas reflects behavioural choices in hawk moths. Scientific Reports, 6(1), 1–10.

Stoddard, M. C., & Hauber, M. E. (2017). Colour, vision and coevolution in avian brood parasitism. Philosophical Transactions of the Royal Society B: Biological Sciences, 372(1724), 20160339.

Strausfeld, N. J. (1989). Beneath the Compound Eye: Neuroanatomical Analysis and Physiological Correlates in the Study of Insect Vision. In Facets of Vision (pp. 317–359). Berlin, Heidelberg: Springer Berlin Heidelberg.

Strausfeld, N. J., Hansen, L., Li, Y., Gomez, R. S., & Ito, K. (1998). Evolution, discovery, and interpretations of arthropod mushroom bodies. Learning & Memory, 5(1), 11–37.

Sulger, E., McAloon, N., Bulova, S. J., Sapp, J., & O’Donnell, S. (2014). Evidence for adaptive brain tissue reduction in obligate social parasites (Polyergus mexicanus) relative to their hosts (Formica fusca). Biological Journal of the Linnean Society, 113(2), 415–422.

Toth, A. L., Varala, K., Henshaw, M. T., Rodriguez-Zas, S. L., Hudson, M. E., & Robinson, G. E. (2010). Brain transcriptomic analysis in paper wasps identifies genes associated with behaviour across social insect lineages. Proceedings of the Royal Society B: Biological Sciences, 277(1691), 2139–2148.

Turillazzi, S., Cervo, R., & Cavallari, I. (1990). Invasion of the nest of Polistes dominulus by the social parasite Sulcopolistes sulcifer (Hymenoptera, Vespidae). Ethology, 84(1), 47–59.

Turillazzi, S., Sledge, M. F., Dani, F. R., Cervo, R., Massolo, A., & Fondelli, L. (2000). Social hackers: integration in the host chemical recognition system by a paper wasp social parasite. Naturwissenschaften, 87(4), 172–176.

Walton, A., Tumulty, J. P., Toth, A. L., & Sheehan, M. J. (2020). Hormonal modulation of reproduction in Polistes fuscatus social wasps: dual functions in both ovary development and sexual receptivity. Journal of Insect Physiology, 120, 103972.

Warton, D. I., Duursma, R. A., Falster, D. S., & Taskinen, S. (2012). smatr 3 – an R package for estimation and inference about allometric lines. Methods in Ecology and Evolution, 3(2), 257–259.

Warton, D. I., Wright, I. J., Falster, D. S., & Westoby, M. (2006). Bivariate line-fitting methods for allometry. Biological Reviews, 81(2), 259–291.

Yang, E. C., Lin, H. C., & Hung, Y. S. (2004). Patterns of chromatic information processing in the lobula of the honeybee, Apis mellifera L. Journal of Insect Physiology, 50(10), 913–925.

Yilmaz, A., Grübel, K., Spaethe, J., & Rössler, W. (2019). Distributed plasticity in ant visual pathways following colour learning. Proceedings of the Royal Society B: Biological Sciences, 286(1896), 20182813.

