## Supplementary material for "Differential investment in visual and olfactory brain regions mirrors the sensory needs of a paper wasp social parasite and its host": Rozanski_etal_Suppl_Material_NeuroAnSocPar

**Supplementary Table 1. Standardized Major Axis regressions to test for allometry in *Polistes dominula* (host) and *Polistes sulcifer* (social parasite).**

Investment in the volume of brain regions that receive and process sensory stimuli was compared to the investment in central brain volume. Scaling of each brain region was calculated with a Grade Shift Index (GSI = $e^{\alpha\ host\ -\ \alpha\ par}$) that compares differences in elevation between the two species (Ott & Rogers, 2010; Sheehan et al., 2019; Stöckl et al., 2016). If GSI >1, there's higher in investment in a specific brain region by the host. If GSI <1, there's higher investment in a specific brain region by the social parasite. The Slope Index (SI) calculates if the allometric scaling of each brain region to the central brain deviates from $\boldsymbol{\beta}$ =1. Each statistical test was implemented as recommended by the SMATR 3 package in R (Warton et al., 2012).

| | **Common Slope** | | **Isometry** | | | **Common Shift** | | **Common Elevation** | | |
|---|---|---|---|---|---|---|---|---|---|---|
| **Brain region** | **Log likelihood** | ***P*** | **Log likelihood** | ***P*** | **SI** | **Wald test** | ***P*** | **Wald test** | ***P*** | **GSI** |
| Whole Brain | 4.926 | 0.026 | | | | | | | | |
| Central Brain | 0.131 | 0.717 | 0.261 | 0.25 | 0.9 | 0.545 | 0.461 | 5.974 | 0.014 | 1.02 |
| Sensory regions | 0.335 | 0.562 | 0.372 | 0.1 | 1.25 | 0.219 | 0.589 | 5.845 | 0.016 | 0.97 |
| Optic Lobe | 0.212 | 0.645 | 0.376 | 0.09 | 1.35 | 0.019 | 0.891 | 18.660 | <0.001 | 0.93 |
| Antennal Lobe | 0.042 | 0.836 | 0.684 | <0.001 | 1.84 | 3.098 | 0.078 | 1.388 | 0.238 | 1.03 |
| Lamina | 0.141 | 0.707 | 0.586 | <0.001 | 1.82 | 0.551 | 0.458 | 28.780 | <0.001 | 0.90 |
| Medulla | 0.123 | 0.726 | 0.327 | 0.15 | 1.28 | 0.015 | 0.903 | 11.220 | <0.001 | 0.95 |
| Lobulla | 3.890 | 0.050 | 0.394 | 0.08 | 1.37 | 0.019 | 0.891 | 6.625 | 0.01 | 0.96 |
| Calyx | 0.168 | 0.682 | 0.674 | <0.001 | 1.66 | 3.851 | 0.042 | 5.891 | 0.012 | 1.04 |
| Collar | 0.177 | 0.674 | 0.722 | <0.001 | 1.8 | 2.943 | 0.086 | 2.089 | 0.148 | 1.03 |
| Lip | 0.768 | 0.087 | 0.7174 | <0.001 | 1.91 | 6.925 | <0.001 | 16.27 | <0.001 | 1.07 |
| Central Complex | 0.848 | 0.357 | 0.521 | 0.02 | 1.75 | 0.009 | 0.357 | 3.547 | 0.045 | 0.93 |

| P ≤ 0.01 | P ≤ 0.05 | P >0.05 |
|---|---|---|

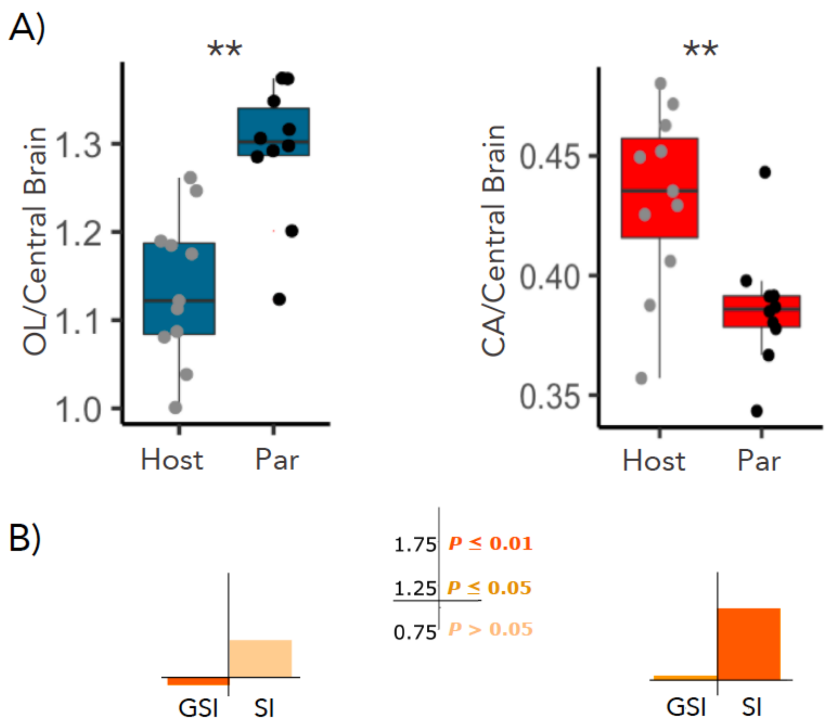


Suppl. Fig 1. A) Volume for normalized optic lobes (OL) and calyx (CA) for hosts and in black for social parasites. Each dot represents an individual. B) The Grade Shift Index (GSI) represents scaling differences between hosts and parasites for these brain regions (GSI > 1, hosts had higher investment in a sensory brain region, and conversely if GSI<1, parasites had higher investment. The Slope Index (SI) represents the deviation of the estimated common allometric slope $\beta$ from 1.Statistical results based on Mann-Whitney U tests (**: $P< 0.001$). Full statistical tests can be found in the results sections and Supplementary Table 1.

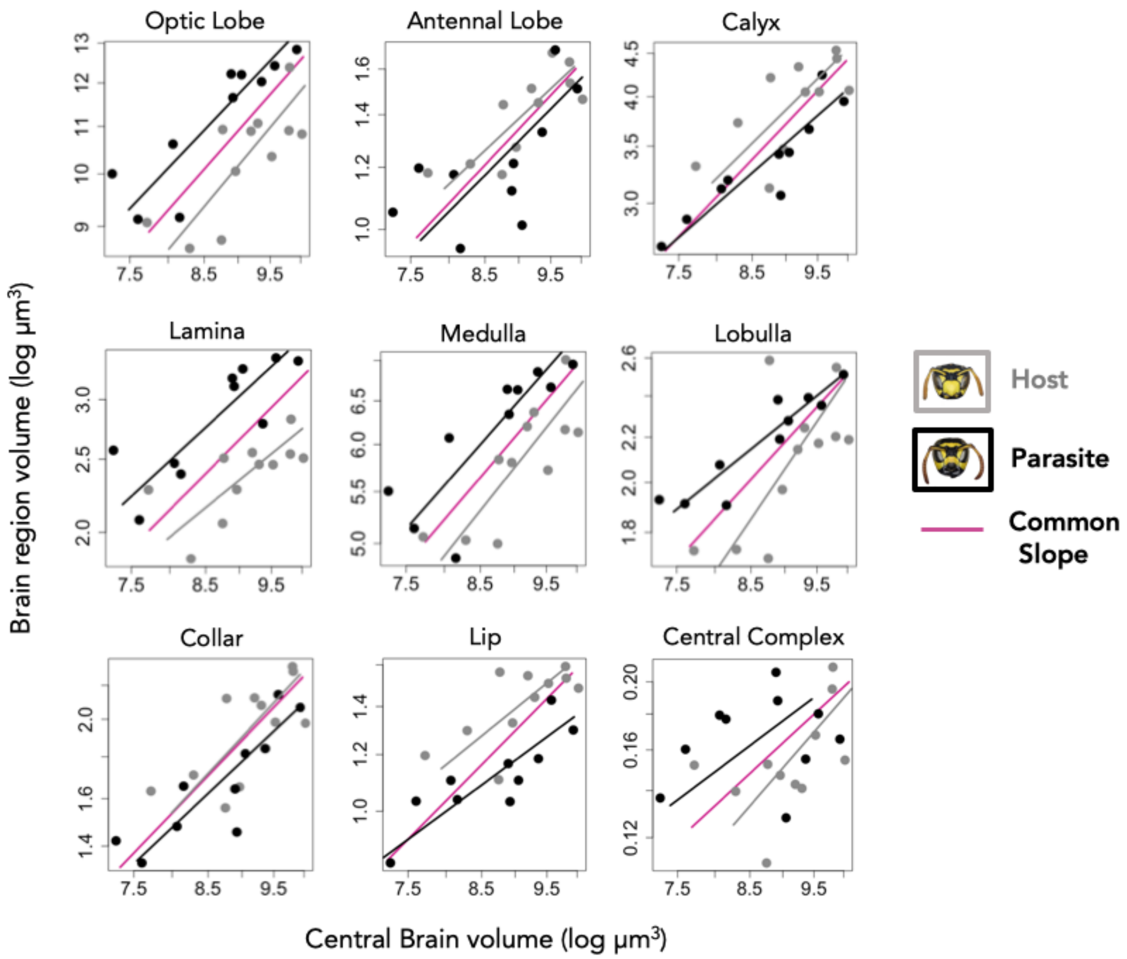


Suppl Fig. 2. **Scaling relationship between the volume of log transformed sensory brain regions and the Central Brain.** The pink line indicates the common allometric slope for the host and parasite species. Grey lines depict the allometric slope for the host and the black line for the parasites. Each dot represents an individual. See Suppl. Table 1 for full Standardized Major Axis Tests and inference for allometric lines.

**Literature cited**

Ott, S. R., & Rogers, S. M. (2010). Gregarious desert locusts have substantially larger brains with altered proportions compared with the solitarious phase. *Proceedings of the Royal Society B: Biological Sciences, 277*(1697), 3087-3096.

Sheehan, Z. B., Kamhi, J. F., Seid, M. A., & Narendra, A. (2019). Differential investment in brain regions for a diurnal and nocturnal lifestyle in Australian Myrmecia ants. *Journal of Comparative Neurology, 527*(7), 1261-1277.

Stöckl, A., Heinze, S., Charalabidis, A., El Jundi, B., Warrant, E., & Kelber, A. (2016). Differential investment in visual and olfactory brain areas reflects behavioural choices in hawk moths. *Scientific Reports, 6*(1), 1-10.

Warton, D. I., Duursma, R. A., Falster, D. S., & Taskinen, S. (2012). smatr 3-an R package for estimation and inference about allometric lines. *Methods in Ecology and Evolution, 3*(2), 257-259.
